# A neuronal circuit driven by GLP-1 in the olfactory bulb regulates insulin secretion

**DOI:** 10.1101/2023.08.28.555086

**Authors:** Mireia Montaner, Jessica Denom, Vincent Simon, Wanqing Jiang, Marie K Holt, Dan Brierley, Claude Rouch, Ewout Foppen, Nadim Kassis, David Jarriault, Dawood Khan, Louise Eygret, Francois Mifsud, David J Hodson, Johannes Broichhagen, Lukas Van Oudenhove, Xavier Fioramonti, Victor Gault, Daniela Cota, Frank Reimann, Fiona M Gribble, Stephanie Migrenne-Li, Stefan Trapp, Hirac Gurden, Christophe Magnan

## Abstract

Glucagon-like peptide 1 (GLP-1) stimulates insulin secretion and holds significant pharmacological potential. Nevertheless, the regulation of energy homeostasis by centrally-produced GLP-1 remains partially understood. Preproglucagon cells, known to release GLP-1, are found in the olfactory bulb (OB). We demonstrate that activating GLP-1 receptors (GLP-1R) in the OB stimulates insulin secretion in response to oral glucose in lean and diet-induced obese mice. This is associated with reduced noradrenaline content in the pancreas and blocked by an α2-adrenergic receptor agonist, highlighting the functional implication of the sympathetic nervous system (SNS). Inhibiting GABAA receptors in the paraventricular nucleus of the hypothalamus (PVN), the control centre of the SNS, abolishes the enhancing effect on insulin secretion induced by OB GLP-1R. Therefore, OB GLP-1-dependent regulation of insulin secretion relies on a relay within the PVN. These findings identify a novel top-down neural mechanism engaged by OB GLP-1 signaling to control insulin secretion via the SNS.

## Introduction

Glucagon like peptide-1 (GLP-1) is an insulinotropic^1,2^ incretin derived from the preproglucagon (PPG) which was first identified in the intestine^3^. Since its discovery in the periphery, the presence of PPG, GLP-1, and its receptor GLP-1R has also been described in the central nervous system (CNS). GLP-1 is notably produced by hindbrain PPG neurons, mainly in the nucleus tractus solitarii (NTS) and the medullary intermediate reticular nucleus (IRT)^4^. In mice, GLP-1R are found in several areas including the circumventricular organs, the amygdala and hypothalamic nuclei in mice^5^. The role of the neural circuit driven by GLP-1 and GLP-1R in the NTS and hypothalamus has been shown to contribute to many aspects of energy balance control^4,6^. Thus, the idea emerged that GLP-1 could be defined as a hormone at the periphery and as a neurotransmitter in the CNS^7–9^.

The olfactory bulb (OB) is another key region in the control of energy balance in rodents^10^ as well as in humans^11–13^ and plays a fundamental role in the appreciation of food palatability. The OB also contains cells expressing PPG and GLP-1R, particularly in the glomerular and granular cell layers^14,15^. Therefore, PPG cells in the OB could be the main source of GLP-1 found in this structure^15^. Additionally, GLP-1R mRNA was also shown to be abundantly expressed in mitral cells (MCs)^14^, the main output neurons of the OB. The presence of GLP-1R in MCs has been recently confirmed by multiple research groups, including our own^5,15,16^. Additionally, *ex vivo* data have shown that MCs increase their action potential firing frequency after the addition of both GLP-1 and the GLP-1R agonist exendin-4^17^. Building on these previous findings, we aimed to provide new insights in the role of OB GLP-1, focusing on its potential pathophysiological implications. Hence, we hypothesized that GLP-1 signaling in the OB could be the trigger for a neural circuitry that regulates energy homeostasis in the context of metabolic disorders.

## Results

### PPG neurons, enzymatic machinery for local processing of GLP-1 and GLP-1R-expressing cells are present in the OB

We first wanted to confirm the presence of neurons expressing PPG and GLP-1R in the OB (Figure 1A). Using transgenic mice expressing the yellow fluorescent protein (YFP) under the control of the glucagon promoter (PPG-YFP mice)^18^ we identified a large population of PPG-neurons in the granular cell layer (GCL) of the OB (Figure 1B and 1C) confirming the presence of PPG cells that we previously evidenced^15,16^.

**Figure 1.**
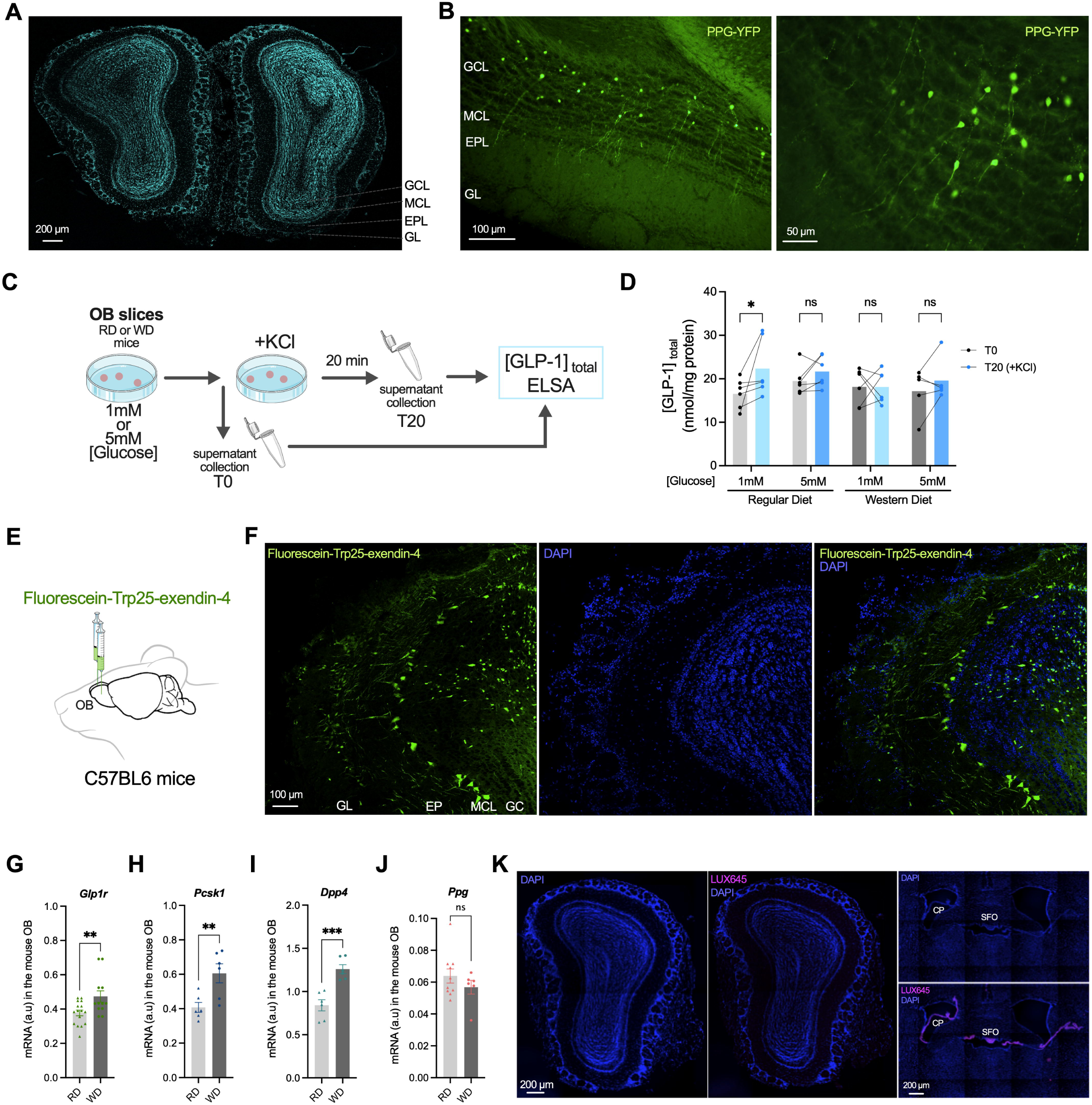
PPG neurons and GLP-1R-expressing neurons are present in the OB. (A) DAPI-stained mouse OB section. GCL, granular cell layer; MCL, mitral cell layer; EPL, external plexiform layer; GL, glomerular layer. Scale bar = 200 µm. (B) Representative photomicrographs of a PPG-YFP mouse OB showing YFP labelling of PPG cells in the GCL. (C) Schematic illustration of the strategy targeting GLP-1 production in OB slices from adult RD or WD mice. (D) Total GLP-1 detection (nmol/mg protein) in OB slices supernatant before (T0) and after (T20) 100 mM KCl addition (n=5-6/group). (E) Strategy to deliver fluorescent Ex4 in the OB of OB-cannulated C57BL6 mice. (F) Photomicrograph of representative coronal OB sections from a fluorescent Ex4-injected mouse. Scale bar = 100 µm. (G-J) RT-qPCR of *Glp1r* (n=16 RD and 12 WD), *Psck1* (n=6), *Dpp4* (n=6) and *Ppg* (n=10 RD and 6 WD) in the OB of RD and WD mice. (K) Photomicrograph of representative OB (left, middle), SFO and surrounding area (right) coronal sections from RD mice IP-injected with LUXendin 645 (LUX645). SFO, Subfornical Organ; CP, Choroid Plexus. Data in (D) are compared using 2-way ANOVA followed by Fisher’s LSD test. Data in (G-J) are compared using unpaired Student’s t-test. Data are given as mean ± SEM. *p <0.05; **p < 0.01; ***p < 0.001; ns, not significant.

To further study the functionality of PPG cells, we performed e*x vivo* experiments and measured the amount of GLP-1 present in the supernatant of OB explants from mice fed a regular diet (RD) or a western diet (WD) leading to obesity and insulin resistance. Explants were exposed to 100mM KCl to depolarize OB neurons (Figure 1C). KCl increased to a small yet significant extent GLP-1 content in the supernatant of OB explants from RD mice exposed to 1mM glucose (Figure 1D), suggesting that GLP-1 in the OB can be secreted by depolarization. However, no such increase in GLP-1 was observed in OB explants from WD mice under the conditions tested (Figure 1D).

We next aimed to assess the location of GLP-1R in the OB. GLP-1R-expressing cells were targeted using a fluorescently labeled Exendin-4 (Fluorescein-Trp25-exendin-4; FLEX) injected locally (Figure 1E). FLEX clearly labelled cells in the mitral cell layer (MCL) and GCL of both RD and WD mice (Figure 1F). Cells with the morphological appearance of MCs were labelled by FLEX in the MCL, as well as scattered cells within the GCL. *Glp1r* mRNA was detected in the OB of both RD and WD-fed mice and found to be significantly increased in the latter (Figure 1G). Interestingly, the enzymatic machinery required for the local processing of GLP-1 was also detected and appeared to be sensitive to diet-induced obesity: mRNAs coding for prohormone-convertase 1/3 (*Pcsk1*) and dipeptidyl peptidase-4 (*Dpp4*) were identified and significantly increased in WD mice compared with RD mice (Figure 1H and 1I). The presence of *Ppg* (Figure 1J) mRNA was also confirmed, but compared to RD mice, their levels remained unchanged in the WD group.

In order to determine whether circulating GLP-1 has access to the OB, LUXendin 645, a fluorescent GLP-1R antagonist based on Ex-9^19^, was administered intraperitoneally (IP). LUXendin 645 stained several GLP-1R-expressing brain structures such as the subfornical organ (Figure K, bottom right), median eminence, arcuate nucleus, organum vasculosum laminae terminalis and area postrema (Figure S1A) but did not label OB GLP-1R (Figure 1K, left and middle). This highlighted that circulating GLP-1 might have restricted, if any, access to the OB. Additionally, icv-injected LUXendin 645 hardly reaches the MCL and GCL of the OB (Figure S1B) and PPG^NTS^ neurons do not project to the OB (Figure S1C). This suggests that neither direct release of GLP-1 in the OB from hindbrain PPG neuron projections, nor volume transmission via release of GLP-1 into the ventricles are likely sources of GLP-1 to activate OB GLP-1Rs.

Taken together, these results indicate that PPG cells located in the OB are the most plausible source of bulbar GLP-1.

### GLP-1R-expressing cells in the OB control glucose homeostasis

To assess the role of the OB GLP-1R-experssing cells in glucose homeostasis and insulin secretion, we first used a pharmacological approach in RD and WD mice (Figure 2A and 2B). Either GLP-1, Exendin-4 (Ex4, GLP-1R agonist), Exendin-9 (Ex9, GLP-1R antagonist) or saline (vehicle) were injected into the OB and an oral glucose tolerance test (OGTT) was performed with measurement of insulinemia (Figure 2C). In RD mice, GLP-1 injection significantly reduced the area under the glucose curve (AUC) during the OGTT (Figure 2D). No significant change was observed after Ex-4 injection (Figure 2E). Ex9 had a deleterious effect on glycemia during the OGTT in RD mice (Figure 2F). Moreover, the injection of GLP-1 and Ex4 into the OB of WD mice tended to normalize the time course of glycemia during OGTT to a value close to that of control RD mice (Figure 2D and 2E). This improvement in glucose tolerance observed in WD mice treated with GLP-1 or Ex4 in the OB was partly due to an increase in insulin secretion during OGTT (Figure 2G and 2H), while insulin sensitivity was also improved following intrabulbar injection of GLP-1 or Ex4 (Figure 2J and 2K). Similar results were obtained during IPGTT: bulbar Ex4 decreased glycaemia (Figure S2A) in RD mice and enhanced glucose tolerance and insulin secretion in obese WD mice (Figure S2A and S2B). Administration of Ex9, on the other hand, exacerbated glucose intolerance (Figure 2F) and insulin resistance (Figure 2L) but did not affect insulin secretion (Figure 2I). Importantly, the effectiveness of this GLP-1 antagonist is in favor of a local release of GLP-1 during OGTT and fits with the GLP-1 release demonstrated in Figure 1C.

**Figure 2.**
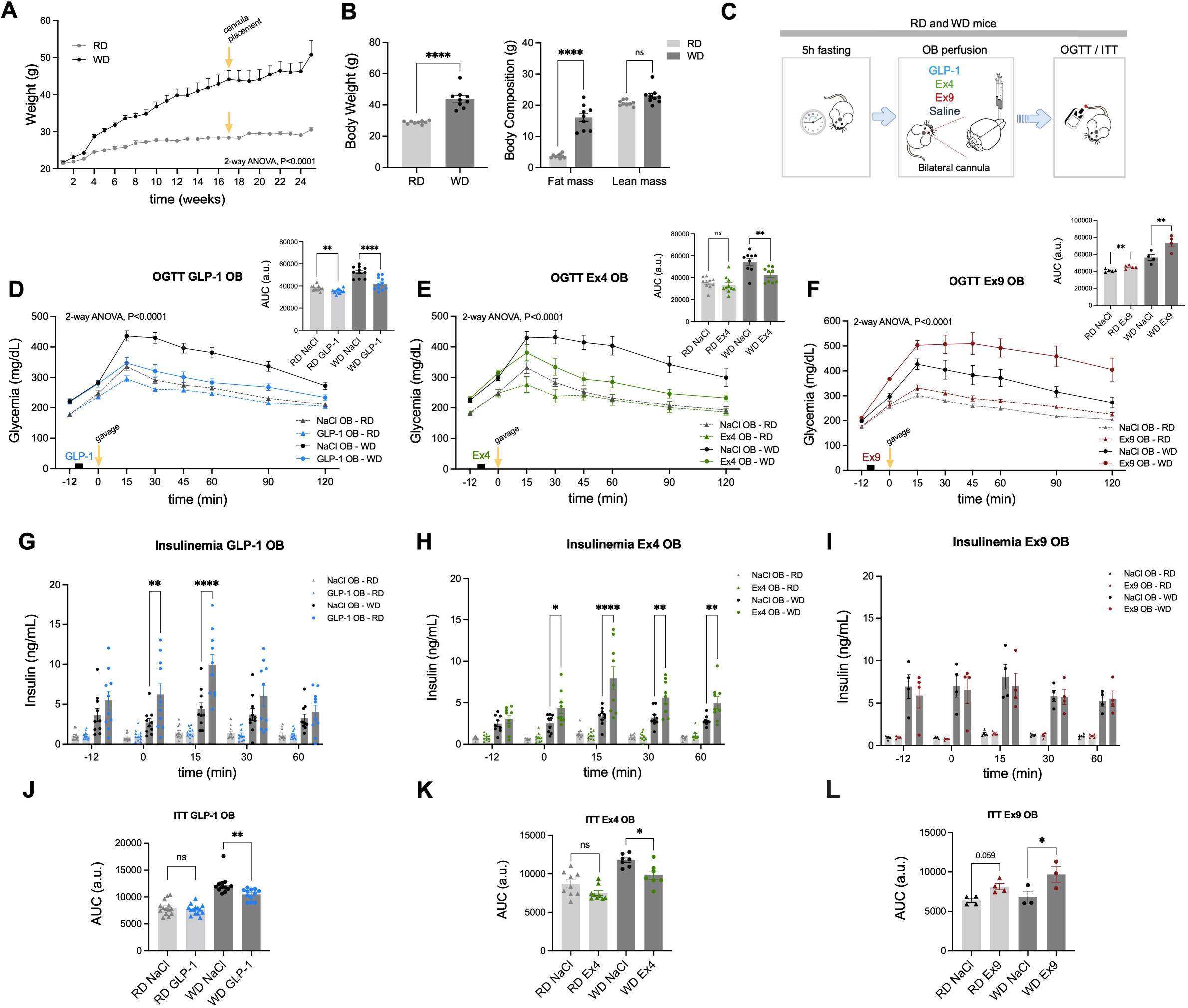
Pharmacological modulation of GLP-1R in the OB regulates glucose homeostasis. (A) 25-week weight gain of age-matched RD and WD mice from week 1 under WD (n=9/diet group). At week 17 mice undergo surgery. (B) Body weight, fat mass and lean mass of OB-cannulated RD and OB-cannulated WD mice at 16 weeks (n=9/diet group). Data are analyzed using Paired Student’s t-test/diet group. (C) Schematic illustration of bilateral OB-injections of GLP-1 (0,5μg/μl), Ex4 (0,5μg/μl) and Ex9 (12,5μg/μl) followed by an OGTT or ITT in OB-cannulated RD and WD mice. (D -F) OGTT tests combined with preceding OB injections of GLP-1 (D; n=11), Ex4 (E; n=10 RD, n=9 WD) or Ex9 (F; n=5 RD, n= 4 WD) in OB-cannulated RD and WD mice after 20 weeks under WD (26-week-old). Black squares on X axis indicate drug delivery before the glucose challenge (T0). Inset above, right: incremental AUC of glycaemia. OGTT curves are analyzed using two-way ANOVA followed by Bonferroni post-hoc test; AUCs are analyzed using Paired Student’s t-test/diet group. (G-I) Plasma insulin levels of OB-cannulated RD and WD mice measured during GLP-1 (G), Ex4 (H) or Ex9 (I) OB-injected OGTT tests (tests in D, E, F respectively). Data are analyzed using two-way ANOVA followed by Bonferroni post-hoc test (n=4-11). (J-L) AUC of ITT tests on mice previously OB-injected with GLP-1 (J; n=14 RD, 12 WD), Ex4 (K; n=9 RD, 7 WD), or Ex9 (L; n=4 RD, n=3 WD). WD mice were fed a WD for 24 weeks. The AUC is measured from baseline (before central perfusions, T-12) until T30 following insulin delivery (T0). Data are analyzed using Paired Student’s t-test/diet group. Data are given as mean ± SEM. *p <0.05; **p < 0.01; ***p < 0.001; ns, not significant.

In addition to pharmacology, we used a chemogenetic approach to drive the activity of *Glp1r*-expressing neurons in the OB. Viral-vector delivery of KORD or hM3Dq receptor-based DREADDs to the OB was performed in *Glp1r*-Cre mice in RD and WD mice respectively (Figure 3A, F3B and F3C). RNAscope was used to visualize *Glp1r* mRNA and validate the hM3Dq chemogenetic model (Figure S3A). We specifically inhibited or activated *Glp1r*-expressing cells in the OB by IP injection of Salvinorin B (SALB) or clozapine N-oxide (CNO), respectively. SALB administration in *Glp1r*^OBKORD^ RD mice had no major effect on glycemia and insulin levels during OGTT compared with controls (Figure 3D and 3E). Conversely, in *Glp1r*^OBhM3Dq^ WD mice, CNO injection induced a significant improvement in glucose tolerance, resulting in a decrease in glycemia AUC (Figure 3F), and a significant increase in insulinemia (Figure 3G). The dose of CNO injected was confirmed to affect neither glycaemia nor insulinemia in WD mice transduced with a control AAV (AAV8-hSyn-DIO-mCherry; Figure S3B and S3C).

**Figure 3.**
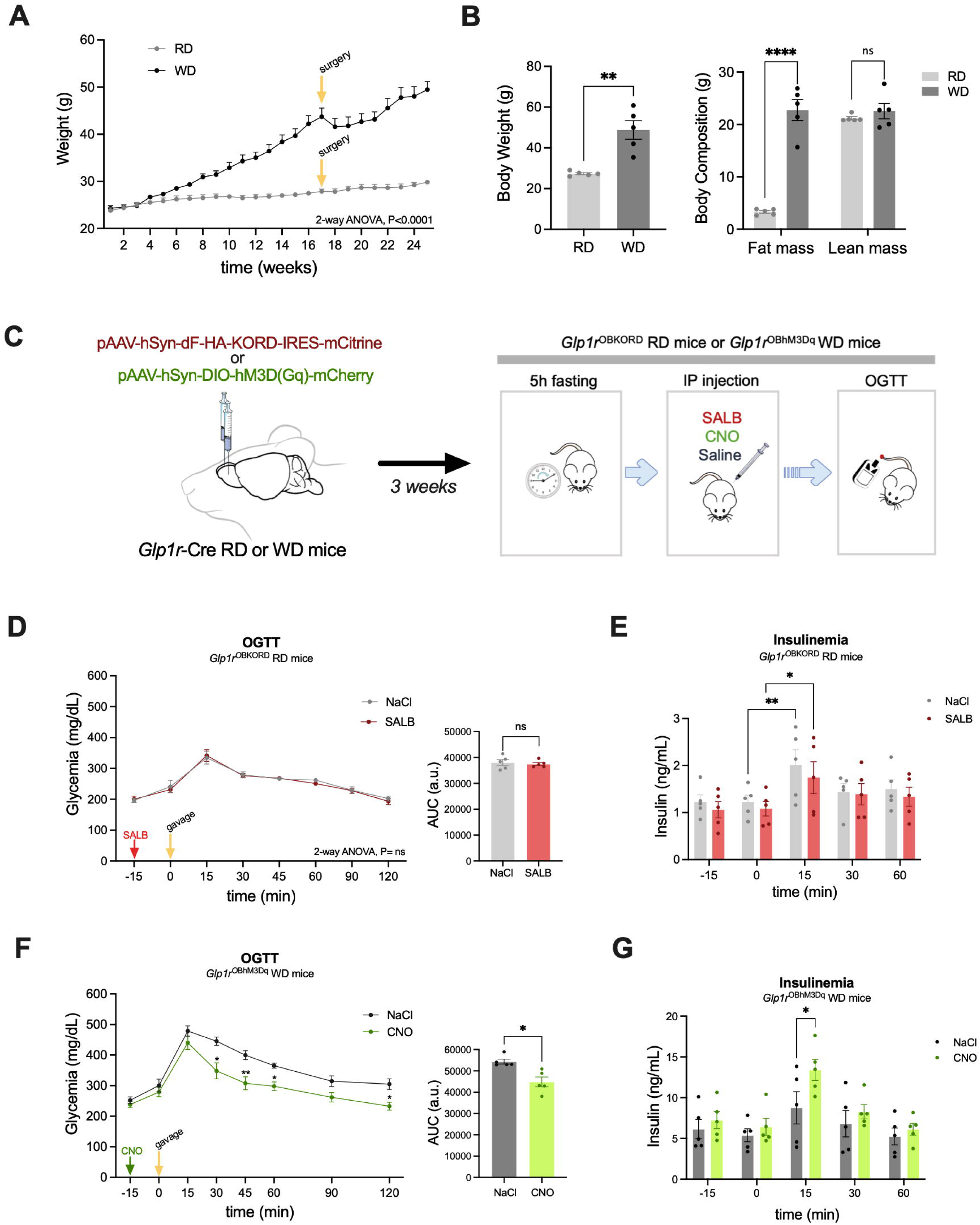
Chemogenetic modulation of GLP-1R+ cells in the OB increases insulin secretion in WD mice. (A) 25-week weight gain of *Glp1r-*Cre RD and WD mice from week 1 under WD (n=5/diet group). At week 17 mice undergo surgery. (B) Body weight, fat mass and lean mass of *Glp1r-*Cre RD and WD mice at 16 weeks under WD diet (n=5/diet group). Body composition (fat and lean mass) was expressed as a percentage of body weight. Data are analyzed using Paired Student’s t-test/diet group. (C) Schematic illustration of viral delivery in the OB of *Glp1r-*Cre RD and WD mice (left). SALB or CNO delivery prior to the onset of the OGTT (right). (D) OGTT preceded by an IP injection of SALB in *Glp1r*^OBKORD^ 26-week-old. The red arrow indicates the SALB injection before the glucose gavage (T0) (left). AUC of glycaemia. OGTT data are analyzed using Repeated Measures (RM) two-way ANOVA followed by Bonferroni post-hoc test and AUCs by using paired Student’s t-test (n=5) (right). (E) Plasma insulin levels of *Glp1r*^OBKORD^ RD mice after SALB IP injection (RM two-way ANOVA followed by Bonferroni post-hoc test; n=5). (F) OGTT tests preceded by an IP injection of CNO in *Glp1r*^OBhM3Dq^ WD mice after 20 weeks under WD (26-week-old, n=5). The green arrow indicates the CNO injection before the glucose gavage (T0) (left). AUC of glycaemia (right). OGTT data are analyzed using RM two-way ANOVA followed by Bonferroni post-hoc test and AUCs by using paired Student’s t-test. (G) Plasma insulin levels of *Glp1r*^OBhM3Dq^ WD mice after CNO IP injection (RM two-way ANOVA followed by Bonferroni post-hoc test; n=5). Data are given as mean ± SEM. *p <0.05; **p < 0.01; ***p < 0.001; ns, not significant.

Moreover, we studied the role of OB *Glp1r*-expressing cells in the control of food intake. In *Glp1r*^OBKORD^ RD mice, SALB injection significantly increased food intake in a fasting-refeeding experiment (Figure S3E). In *Glp1r*^OBhM3Dq^ WD mice fed a both a regular and a high fat/high sucrose diet, a decrease in food intake was observed after CNO injection (Figure S3F and S3G). With this experiment, we demonstrated that modulating OB *Glp1r*-expressing could modulate food intake.

Overall, using pharmacological and chemogenetic approaches, we provide evidence that the modulation of GLP-1/GLP-1R activity levels in the OB contributes to the maintenance of energy homeostasis and that activation of this ligand/receptor pair can improve metabolic parameters deregulated in the context of obesity.

### OB GLP-1R stimulates insulin secretion by inhibiting the sympathetic nervous system activity

We hypothesized that the increase in insulin secretion during OGTT could be due to changes in autonomic nervous system activity. More precisely, either an increase in parasympathetic nervous system (PNS) activity or a decrease in sympathetic nervous system (SNS) activity can lead to a stimulation of insulin secretion^20^ (Figure S4A). To explore the role of the PNS, we performed subdiaphragmatic vagotomy to cut parasympathetic efferents, including those projecting to pancreatic β-cells (Figure S4B; validated by Fluorogold IP injection, Figure S4E). While RD mice insulinemia remained intact (Figure S4C), the improvement of insulin secretion induced by Ex-4 administration in the OB of obese WD mice was still present after vagotomy (Figure S4D). This indicates that the PNS was not mediating the beneficial effects of Ex-4 administration in the OB.

We then tested the alternative hypothesis that a decrease in SNS activity could account for the increase in insulin secretion following the activation of OB GLP-1Rs. We first measured pancreatic noradrenaline levels following injection of Ex-4 into the OB (Figure 4A). We observed a drop in noradrenaline levels, which could be indicative of a drop in sympathetic tone^21^. To validate this hypothesis of reduced sympathetic tone as a relay for the effect of GLP-1 in the OB, we performed OGTT with or without Ex4 administration in the OB and with or without prior IP injection of an α_2_-adrenergic receptor agonist, UK14304^22^ (Figure 4B). The α_2_-adrenergic pathway is the main inhibitor of insulin secretion induced by SNS activity^20^. Our aim was therefore to activate α_2_ adrenergic receptors on pancreatic β-cells to counteract a decrease in SNS activity triggered by OB GLP-1R. Under these conditions, the beneficial effect of GLP-1 in the OB on both improvement of glucose tolerance and insulin secretion was effectively lost (Figure 4C and 4D). Together, these results suggest that the activation of OB GLP-1R leads to an increase in insulin secretion through the inhibition of sympathetic nervous system (SNS) activity.

**Figure 4.**
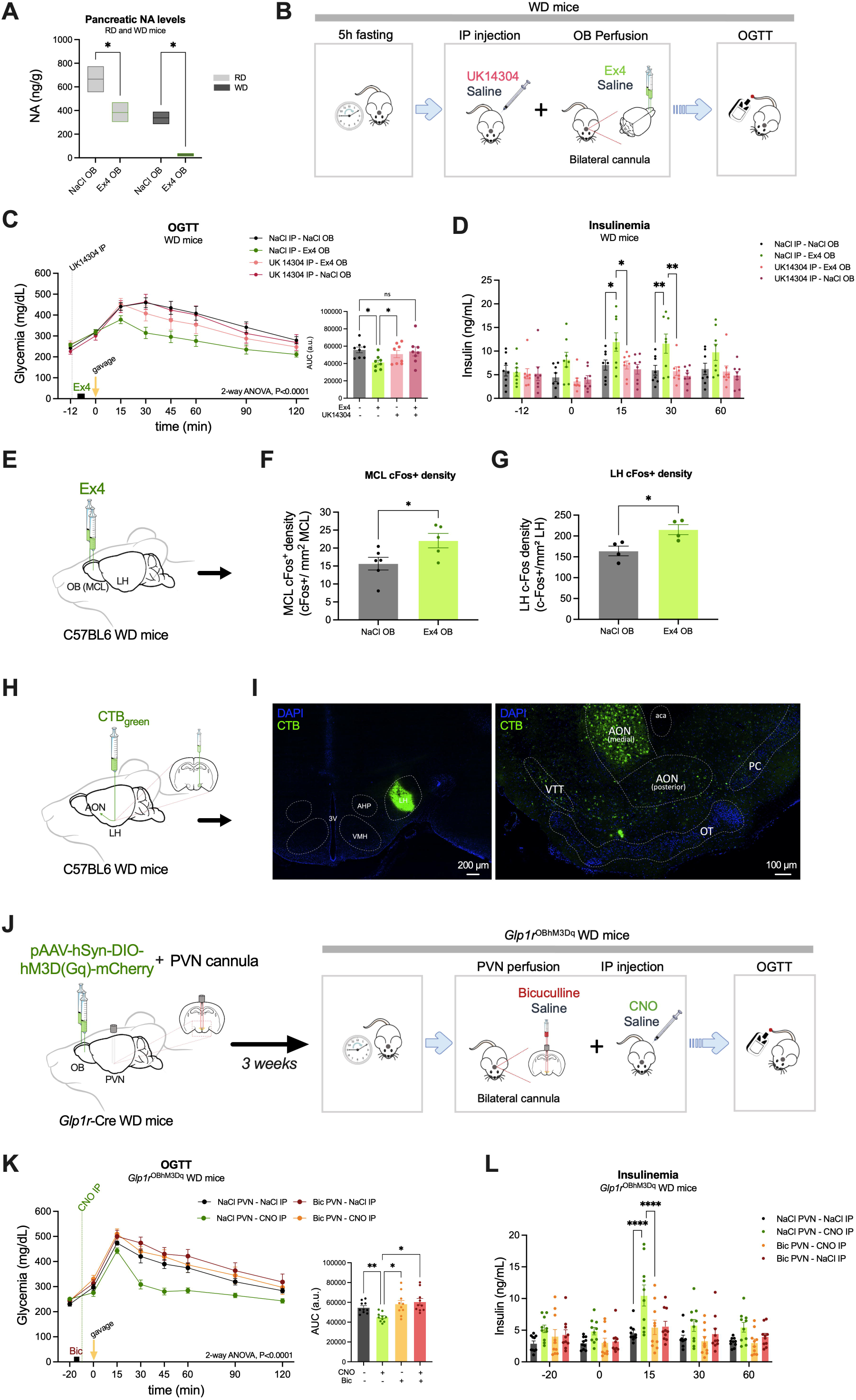
The stimulatory effect of GLP-1 in the OB on insulin secretion is mediated by a decrease in sympathetic nervous system activity. (A) Pancreatic NA levels of RD and WD mice after Ex4 injection in the OB (n=2-3/group). (B) Schematic illustration of bilateral OB injections of Ex4 combined with IP administration of UK14304 followed by an OGTT in OB-cannulated WD and OB-cannulated RD mice. (C) OGTT tests preceded by an IP injection of UK14304 and an OB injection of Ex4 in OB-cannulated WD after 20 weeks under WD (n=8) (left). AUC of glycaemia (right). OGTT data are analyzed using RM two-way ANOVA followed by Bonferroni post-hoc test and AUCs by using paired Student’s t-test. (D) Plasma insulin levels of OB-cannulated WD during OGTT tests preceded with OB-injected Ex4 and IP injected UK14304 (RM two-way ANOVA followed by Bonferroni post-hoc test; n=8). (E) Strategy to deliver Ex4 in the OB of OB-cannulated C57BL6 mice. (F) Representative photomicrographs of cFos immunoreactivity and automated cell counting in the MCL using CellProfiler software (left). cFos density in the MCL after Ex4 or saline injection in the OB in WD mice (right). Data are expressed as cFos+ nuclei/mean MCL area of all mice. Data are analyzed using an unpaired Student’s t-test (n=5). (G) cFos density in the LH after Ex4 or saline injection in the OB in WD mice. Data are expressed as cFos+ nuclei/mean LH area of all mice. Data are analyzed using an unpaired Student’s t-test (n=4). (H) Schematic illustration of unilateral Alexa 488-conjugated cholera toxin B subunit (CTB_green_) delivery in the LH for retrograde tracing of the primary olfactory cortex. (I) Coronal sections of the LH after unilateral injection of CTB_green_ (left). CTB staining in the primary olfactory cortex (right). LH, lateral hypothalamus; AHP, anterior hypothalamus; VMH, ventromedial hypothalamus; 3V, third ventricle; AON, anterior olfactory nucleus; VTT, ventral tenia tecta; VON, ventral olfactory nucleus; PC, piriform cortex; OT, olfactory tubercle. Structure boundaries were drawn based on Franklin and Paxinos mouse brain atlas. (J) Schematic illustration of viral delivery in the OB and bilateral cannula placement in the PVN of *Glp1r*-Cre WD mice followed by OGTTs. (K) OGTT tests preceded with an IP injection of CNO in *Glp1r*^OBhM3Dq^ WD PVN-cannulated mice after 20 weeks under WD (26-week old; n=10) (left). The black square and the green line indicate Bicuculline and CNO delivery, respectively, before the glucose gavage (T0). AUC of glycaemia (right). OGTT data are analyzed using RM two-way ANOVA followed by Bonferroni post-hoc test and AUCs by using RM one-way ANOVA followed by Bonferroni post-hoc test. (L) Plasma insulin levels of *Glp1r*^OBhM3Dq^ WD PVN-cannulated mice after CNO IP injection (RM two-way ANOVA followed by Bonferroni post-hoc test, n=10). Data are given as mean ± SEM. *p <0.05; **p < 0.01; ***p < 0.001; ns, not significant.

### The OB GLP-1R-induced decrease in sympathetic nerve activity is GABA dependent and relays in the PVN

We aimed to identify the mechanisms leading to the decrease in SNS activity after activation of the GLP-1/GLP-1R neuronal circuit in the OB, which was confirmed by an increased cFos expression in the MCL after local Ex4 injection (Figure 4E and 4F). These results were expected given the reported increased excitability of mitral cells, due to an increase in the firing frequency, after GLP-1 or Ex4 addition in OB slices^17^. Pre-autonomic neurons are located in the PVN^23^ and are partly controlled by inhibitory GABA projections (through GABA_A_ receptors) from the lateral hypothalamus (LH)^24–26^.

In support of a multisynaptic OB to LH pathway, retrograde tracing (Figure 4H) revealed neurons in the anterior olfactory nucleus (AON), a major output of the OB, that project to the LH (Figure 4I). Furthermore, injection of Ex-4 into the OB elicited an increase in cFos expression in LH neurons (Fig 4G), suggesting that the OB GLP-1 pathway drives activity in the LH.

Considering this evidence, we hypothesized that activation of the GLP-1 pathway in the OB activates GABAergic neurons in the LH, which inhibit PVN neurons, reducing their stimulation of preganglionic sympathetic neurons.

To test whether a GABA-dependent decrease in the sympathetic tone could explain the effect on insulin secretion mediated by GLP-1R activation in the OB, we performed OGTTs on *Glp1r*^OBhM3Dq^ WD mice after chemogenetic stimulation of OB *Glp1r*-expressing neurons under blockade of GABA_A_ receptors in the PVN (by using the GABA_A_ receptor antagonist Bicuculline, Figure 4J). Under these conditions, the stimulatory effect of the GLP-1R pathway in the OB on insulin secretion and the consequent improvement in glucose tolerance were lost (Figure 4K and 4L). We concluded that a GABAergic inhibition in the PVN, driven by OB *Glp1r*-expressing neurons, could decrease sympathetic nerve activity to the pancreas, increasing insulin release in WD mice.

## Discussion

Here we have shown for the first time that GLP-1/GLP-1R in the OB is involved in the central control of glucose homeostasis. In obese and insulin-resistant mice, activation of this system improved glucose tolerance by increasing insulin secretion and action. We proved that increased insulin secretion was due to a decreased SNS activity through a GABA-dependent transmission in the PVN.

Previous studies in rodents have indicated that OB MCs are glucosensitive^27,28^, and chronic activation of these cells in diet-induced obese mice improved glucose tolerance^29^. However, the latter study did not quantify insulin levels, and the central and bulbar mechanisms responsible for the improved glucose tolerance were not elucidated. In our experiments, we showed that local Ex4 injection led to increased neuronal activity in the OB. This activation triggered a series of events involving a specific network, including the hypothalamus, resulting in enhanced insulin secretion due to relief from SNS-inhibition and a metabolic improvement in obese mice. This work parallels our recent study showing that GLP-1 from the OB also controls odor-evoked cephalic phase of insulin secretion^16^. Thus, this neuronal circuit driven by GLP-1 released within the OB could be involved in the physiological control of insulin secretion both in the pre-prandial and post-prandial states. Interestingly, because the activation of OB GLP-1R is efficient to decrease glucose in both RD and WD mice, we demonstrate the absence of GLP-1 resistance in the OB of obese mice. This is in line with the absence of GLP-1 resistance despite the presence of insulin resistance shown in obese humans^30^.

Our study deciphered the connections and mechanisms underlying the action of GLP-1 in the OB and its effect on insulin secretion. We have shown that the stimulatory effect of GLP-1 in the OB on insulin secretion was relayed via the activation of GABA signaling in the PVN and finally decreased SNS activity onto pancreatic β cells. In addition, we have observed increased neuronal activity in the LH upon OB GLP-1R stimulation and the existence of an anatomical AON-LH connection. This is in line with existing anatomo-functional data between OB and hypothalamus. Indeed, these structures share common neuroanatomical pathways and are reciprocally connected^11^. Electrical stimulation of the OB induces an electrophysiological response in the LH and the PVN^31,32^. Additionally, many of the structures to which the OB projects via the MCs, such as the AON, the olfactory tubercle and the piriform cortex, also project to the LH^32–34^. Direct projections of MCs from the OB to the arcuate nucleus (ARC) also exist^35^ and food odors can rapidly inhibit AgRP neurons and excite POMC ones^36,37^. Noteworthy, in the OB, GLP-1 acts on GLP-1R mainly expressed by MCs which also express receptors for leptin, insulin and ghrelin^38^, providing an additional link to the role of these cells in the control of metabolism. In the opposite direction, afferents from various hypothalamic nuclei project to the olfactory system^39,40^. Particularly, the OB is the direct target of orexinergic fibers originating from the LH^41,42^. In addition, some neurons from the PVN project to the olfactory tubercle^43^, a direct target of the OB for the hedonic regulation of smell. Thus, close links exist between these two brain areas involved in the control of energy balance. Our data add a new piece to the puzzle by showing that the GLP-1/GLP-1R signaling in the OB modulates hypothalamic circuits to control pancreatic β-cell function.

In our study, the OB GLP-1/GLP-1R neuronal circuit appears to control insulin secretion via the SNS. A connection between the olfactory system and the periphery through the SNS has previously been proposed by Riera et al.^10^. The authors suggested that a reduction in the olfactory input stimulates SNS activity, resulting in an increase of circulating noradrenaline^10^. Our results mirror these data since we showed that GLP-1R activation in OB decreased noradrenaline content in the pancreas, probably as a result of reduced SNS tone, and this led to improved metabolic outcomes (i.e. insulin secretion and sensitivity). Consistent with the proposed mechanism of action, systemic application of an α_2_-adrenergic receptor agonist abolished the stimulatory effect of GLP-1 in the OB on insulin secretion.

Another important finding of our study was that the inhibitory effect of OB GLP-1 signaling on the SNS was dependent on a GABAergic relay in the PVN. The PVN provides coordination in autonomic and neuroendocrine activities for proper regulation of metabolism. It contains pre-autonomic neurons that control pancreatic SNS activity and insulin secretion^44^. Intra-PVN regions such as parvocellular or mediocellular subdivisions contain neurons projecting to the spinal cord and the medulla. These neurons play a crucial role in the regulation of SNS activity and are regulated by a GABAergic inhibitory input via GABA_A_ receptors (GABA_A_R)^45^. GABAergic neurons from the LH project to GABA_A_R-expressing neurons, as supported by previous studies^26^. Furthermore, our data revealed that the activation of the GLP-1R in the OB results in increased neuronal activity in the LH, as evidenced by cFos labeling. This suggests that the GABAergic innervation of the PVN may originate from the LH, which, in turn, receives projections from the AON, as shown in our study by employing CTB retrograde tracing from the LH. Altogether our results support the existence of a OB → AON → LH → PVN connection. To further describe the mechanism in the PVN relay necessary for the insulin regulation by the OB GLP-1 system, we administered a GABA_A_ receptor antagonist (Bicuculline) locally in the PVN. The results showed that the increased insulin secretion following chemogenetic activation of *Glp1r-*expressing cells is controlled by a GABAergic inhibition of GABA_A_R-expressing neurons in the PVN. This is consistent with previous studies providing evidence for hypothalamic control of sympathetic pre-autonomic neurons in the PVN^25^ through the use of the same GABA_A_ receptor antagonist^46^.

It is of course important to place all these data in a physiological context. Our study showed that there is a local production of GLP-1 in the OB, presumably independent of peripheral GLP-1. This system could be stimulated in the first minutes before a meal and would be involved in foraging and the cephalic phase of insulin secretion, as we have recently shown^16^. During mealtime, the network presented in this study driven by OB GLP-1R, may contribute to a decrease in SNS activity and thus participate in the increase of glucose-induced insulin secretion, in concert with systemic GLP-1 and the GLP-1 neuronal circuitry of the brainstem and hypothalamus under normal physiological circumstances. However, interestingly, it also becomes activated during periods of nutritional stress such as obesity and insulin resistance, as we showed in our data from obese mice. This activation contributes to an improvement of glucose homeostasis, increasing insulin secretion and regulating blood glucose levels.

Additionally, we observed an upregulation of the required enzymatic machinery for local processing of GLP-1 including *Glp1r, Pcsk1* and *Dpp4* mRNAs in the OB of obese mice compared to lean mice. The upregulation of *Glp1r* mRNA may potentially compensate for the lack of increase in GLP-1 release in obese mice OB (Figure 1D). These data provide valuable insights into the adaptive mechanisms operating within the OB in response to metabolic diseases^11^. Because *Glp1r* mRNA is also expressed in the human OB^47^, it becomes intriguing to explore in future studies whether intranasal administration of GLP-1 agonists^48^ could potentially exert their effects by acting in the OB (a site of persistent sensitivity to GLP-1 in obesity according to our data).

Altogether, this work is to be put in perspective with the known role of the OB as a metabolic sensor able to detect variations of insulin and glucose^28^. Olfaction has been recognized as having significant impact on the control of energy balance^11^ and olfactory disorders have been shown to influence eating behavior and whole body metabolism^16,29^. Based on our results, we propose that the GLP-1/GLP-1R system in the OB constitutes a novel neuronal pathway involved in regulating energy homeostasis. This finding holds promise as a potential new target for addressing obesity and type 2 diabetes mellitus.

### Limitations of the study

First, we showed the involvement of GLP-1/GLP-1R in the OB in the control of insulin and glucose only in male obese mice. Considering the distinct metabolic responses of female mice during hypercaloric diet characterized by a strong resistance to weight gain and glucose dysregulation^29^), they are an interesting subject for our follow up studies. Moreover, while our findings have not yet shed light on the functional connection between the OB, the AON and the LH, this presents a captivating area for further studies. The AON has recently been identified as an olfactory hub with significant functions in olfactory gating and memory^49^. Hence, it offers an exciting avenue to understand the link between the olfactory system and the LH in the regulation of energy homeostasis^32,50^. Finally there is not human evidence for a similar OB GLP-1-based circuitry. Human experimental medicine trials with intranasal GLP-1R agonists are in their infancy^48^ and will have to be further developed to understand the translational relevance of the findings presented in this study.

## Author contributions

Conceptualization: C.M., H.G., S.M.L., M.M.; Methodology: C.M., H.G, M.M., S.M.L., S.T. Investigation; M.M., J.D., H.G., D.J., V.S., M.H., D.B., D.H., J.B., W.J., C.R., E.F., N.K., D.K. Writing-original draft: C.M., M.M., H.G. Writing-review and editing: L.V.O., X.F., V.G., M.K.H., D.C., F.R., F.M.G., S.T.

## Acknowledgments

We thank Dr Serge Luquet, Giuseppe Gangarossa and Claire Martin for their advice and helpful discussions. The help of Zahra Boudra is also acknowledged. We thank the technical platform Functional and Physiological Exploration platform (FPE) of the Université de Paris (BFA-UMR 8251), the animal core facility Buffon of the Université de Paris/Institut Jacques Monod for animal experiments.

We also thank the core imaging facility of Institut Jacques Monod (ImagoSeine facility) for help in image acquisition and processing. The microscopy for LH cFos analysis was done in the Bordeaux Imaging Center, a service unit of the CNRS-INSERM and Bordeaux University, member of the national infrastructure France BioImaging supported by the French National Research Agency (ANR-10-INBS-04). The help of Sébastien Marais is acknowledged.

This work was supported by Research grant of the French society for study of diabetes (SFD-017-2019 to CM and HG, the Medical Research Council UK (MR/N02589X/1 to ST), a British Heart Foundation Postdoctoral Fellowship (FS/IPBSRF/20/27001 to MKH) and a grant from the European Foundation for the Study of Diabetes Germany (Merck Sharpe Dohme grant to ST). Additional support was obtained from the Cities Partnership program UCL to ST, CM and HG. MM is supported by a “Université Paris Cité IdEx” PhD fellowship and a EUR GENE, G.E.N.E. Graduate School fellowship. WJ is supported by a UCL ORS scholarship and a CSC scholarship from the Chinese Government. We also acknowledge INSERM (to D.C.) and Agence Nationale de la Recherche (ANR-21-CE14-0018 to D.C.). DJH was supported by MRC (MR/S025618/1) and Diabetes UK (17/0005681) Project Grants, as well as a UKRI ERC Frontier Research Guarantee Grant (EP/X026833/1).

## Methods

### Animals

All animal procedures were performed with approval of the Buffon Ethics Committee (CEEA40), under agreement no. B751317, conforming with the French Ministry of Research and European legislations. Experiments conducted in the United Kingdom were performed in accordance with the UK Animals (Scientific Procedures) Act 1986, and experimental protocols were approved by the UCL Animal Welfare and Ethical Review Body (Bloomsbury Campus). All mice were individually housed in ventilated cages in a temperature (22 ± 2°C) and humidity-controlled room unless otherwise stated. Mice were maintained on a 12-h light/dark cycle with free access to food and water. All mice were accustomed to daily manipulation. All experiments were performed in male mice between 26 and 36 weeks of age.

### Mouse models

A diet-induced obesity model was generated on C57BL/6J male mice (Janvier Labs, Le Genest Saint Isle, France) fed both a mixed high fat/high sucrose (western) diet (HF230 diet, 5317 kcal/kg, Safe, Augy, France) and a regular chow diet (Complete maintenance diet A04, 2791 kcal/kg, Safe, Augy, France) *ad libitum* throughout the whole study from 6 weeks of age.

For selective Cre-dependent viral targeting, we used *Glp1r*-Cre mice (C57BL/6 background)^51^. From 6 week of age, they were fed the same way as DIO mice. All experiments run on both models were done after 26 to 44 weeks of western diet (WD). Age-matched control groups were fed a regular chow diet (RD) *ad libitum*.

PPG-YFP^17^ mice were bred in-house (UCL, London) on a C57BL/6 J background and fed a regular chow diet (Teklad 2018 or 7912, Envigo) *ad libitum*.

### Body composition measurements

Body weight was measured weekly throughout the study, between 09:00 and 10:00, unless otherwise stated. Body mass composition (amount of lean and fat mass) was assessed using an Echo Medical Systems EchoMRI 100 (EchoMRI, Houston, TX, USA) at week 16. RD mice weighting more than 40g were excluded from all the procedures assessed in this work.

### Bilateral OB injections

After 17 weeks under WD (23 weeks of age), mice received an intraperitoneal injection (IP, 10 mg/kg) of Buprécare® (Buprenorphine 0.3 mg) diluted 1/100 in NaCl 0.9% (saline). Mice were anesthetized with isoflurane (Isoflurin 1000 mg/ml, Axience, France) and positioned in a stereotaxic frame (Phymep – Model 940 Small Animals). A double stainless steel guide cannula (26 gauge in diameter, Phymep, France) was implanted into the OB through stereotactic procedures (A/P: +4.8 mm from bregma, M/L: ±1 mm and D/V: -1.7 mm from brain surface) after exposing and properly scrubbing the skull of the animal. To fix the guide cannula onto the skull, plastic screws and dental cement (Super-Bond Universal Kit, Sun Medical and Unifast Trad) were used. A dummy was placed in the guide cannula to assure its patency. Finally, a reversible cap was screwed in the guide cannula to protect its tip. Mice received an IP injection (10 mg/kg) of Ketofen® (Ketoprofen 100 mg) diluted 1/100 in saline and were placed on a heated pad until recovery from anesthesia. Their body temperature was maintained at 37°C throughout the entire surgery. Mice were single housed 1 week before surgery and metabolic tests started after a recovery period of 3 weeks. Mice were acclimated to manipulation for 2 weeks before the tests. On the test day, acute injections in the OB of Exendin-4 (Ex4; Sigma-Aldrich; 0.5μg/μL), Fluorescein-Trp25-Exendin-4 (FLEX; Eurogentec; 0.5μg/μL), Exendin-9 (Ex9; Sigma-Aldrich; 12,5μg/μL or 25 μg/μL), GLP-1 (Sigma-Aldrich; 0.5μg/μL) or saline, for 4 minutes (0,125μL/min), bilaterally, were performed 12 minutes before the beginning of the metabolic tests. No DPP-4 inhibitors were combined with GLP-1 injections.

### Vagotomy

After 26 weeks under WD (32 weeks of age), 6 cannulated WD mice and 6 cannulated RD mice were submitted to a bilateral sub-diaphragmatic vagotomy. Before surgery and for 3 days post-surgery, mice were fed with jelly food (DietGel Boost, Clear H_2_O) to avoid solid food in the digestive tract. Mice received an IP injection (10 mg/kg) of Buprécare® (Buprenorphine 0.3 mg; diluted 1/100 in saline) and were anaesthetized with isoflurane during surgery. Their body temperature was maintained at 37°C throughout the entire surgery. Vagus nerve branches (right and left) were cautiously isolated along the esophagus using a binocular loupe and sectioned in vagotomized mice or left integral in sham ones. Mice recovered for at least 3 weeks post-surgery before undergoing metabolic tests.

### Verification of vagotomy with FluoroGold

To confirm vagotomy, each mouse received an IP injection of FluoroGold (0.8 mg/0.4 ml saline; Sigma-Aldrich). One week after the injection, all animals were transcardially perfused with phosphate-buffered saline (PBS) 1X (pH 7.4) followed by ice-cold 4% paraformaldehyde (PFA; pH 7.4). The brains were removed and post-fixed in 4% PFA overnight, and cryoprotected in 30% sucrose until the tissue sank. After cryoprotection, the brainstem was separated and cut on a cryostat (Leica 1800) into 18 μm coronal sections, which were mounted on microscope slides (SuperFrost Ultra Plus™ GOLD), for later imaging using fluorescence microscopy. Each brainstem section was examined for FluoroGold label in the DMNX. The presence of fluorescent label in the DMNX was accepted as a marker of incomplete vagotomy. Only animals with complete or no vagotomy (for shams) were included in the study.

### Viral injections in the OB

After 17 weeks under WD (23 weeks of age), 11 *Glp1r*-Cre RD and 16 *Glp1r*-Cre WD mice were anesthetized with isoflurane as described above and placed on a stereotaxic frame. Bilateral injections of AAV vectors encoding for stimulatory (AAV8-hSyn-DIO-hM3Dq-mCherry) or inhibitory (AAV8-hSyn-dF-HA-KORD-IRES-mCitrine) Designer Receptors Exclusively Activated by Designer Drugs (DREADDs) were then performed in the OB (A/P: +4.8 mm from bregma, M/L: ±1 mm and D/V: -1.7 mm from brain surface) at a rate of 0.125 μL/min for 4 min, on *Glp1r*-Cre RD and WD mice. The incision was sutured with Coated VICRYL suture (Ethicon). Mice were placed in cages on a heated pad until they recovered from anesthesia. Mice were single housed 1 week before surgery and metabolic tests started after a recovery period of 3 weeks. Experimental groups were called *Glp1r*^OBKORD^ RD and *Glp1r*^OBhM3Dq^ WD respectively.

### Bilateral PVN injections

PVN cannulation was performed on 10 *glp1r*-Cre mice previously injected with AAV8-hSyn-DIO-hM3Dq-mCherry on the same surgery session. A double stainless steel guide cannula (26 gauge in diameter, Phymep, France) was implanted into the PVN through stereotactic procedures (A/P: -0.9 mm from bregma, M/L: ±0.3 mm and D/V: -4.8 mm from dura mater) as described above. PVN-cannulated *Glp1r*^OBhM3Dq^ mice were PVN-infused with Bicuculline Methiodide (Sigma-Aldrich; 0.25μg/μL) or saline (bilaterally, for 2 minutes at a rate of 0.05μL/min).

### Oral glucose tolerance test

Mice were submitted to habituations 1-2 weeks before starting metabolic tests.

Oral glucose tolerance test (OGTT) was performed, unless otherwise stated, 12 minutes after pharmacological injections or 15 minutes after chemogenetic manipulation on mice fasted for 5 hours. Three experimental groups underwent OGTTs: i) WD and RD cannulated mice OB-infused with GLP-1, Ex4 or Ex9 [combined with IP injections of either Ex4 (3 µg/kg) or the α_2_-AR agonist UK14304 (Bio-Techne; 1μg/kg) when stated]; ii) Vagotomized WD mice OB-infused with Ex4; iii) *Glp1r*^OBKORD^ RD and *Glp1r*^OBhM3Dq^ WD mice in which OB *Glp1r*-expressing neurons were chemogenetically activated by IP injections of Salvinorin B (SALB; Sigma-Aldrich; 10mg/kg) or Clozapine N-oxide (CNO; Sigma-Aldrich; 3 mg/kg) respectively (combined with acute PVN injections of Bicuculline Methiodide 3 min prior to CNO administration when stated). A glucose solution (1g/kg) was administrated by oral gavage. Blood was collected from the tail vein for glucose monitoring (Glucofix Tech, Menarini Diagnostics; mg/dL) and blood sampling. Blood cells were removed from plasma by centrifugation in order to assay plasma insulin (ng/mL) and C-peptide (ng/mL) with a wide-range Ultra Sensitive Mouse Insulin ELISA Kit (catalog no. 90080; Crystal Chem, Inc.) and Mouse C-Peptide ELISA Kit (catalog no. 90050; Crystal Chem, Inc.), respectively. IP glucose tolerance tests (IPGTT) were performed the same way, starting with an IP administration of glucose at 1g/kg.

For all glucose tolerance tests, mice receiving a control treatment were administered an equivalent volume of saline. Comparison of treatments was done on a within-subject level (each mouse was its own control) and at least 1 week elapsed between each testing condition (drug or control treatment). Total area under the curve (AUC) of glycaemia was measured from baseline (before central injections) to the end of the glucose tolerance tests, at time = 120 minutes (T120).

### Insulin tolerance test

Insulin tolerance tests (ITT) were performed after 5 hours of fasting in cannulated RD and WD mice. Pharmacological injections of GLP-1, Ex4 or Ex9 were performed in the OB. Mice were given an IP injection of insulin (0.75 U/kg of body weight; Novo Nordisk), and glycemia (mg/dL) was monitored following the same protocol as was used for glucose tolerance tests. For all ITTs, each mouse was its own control. Change the way to say it: The AUC of glycaemia was measured from baseline (before central perfusions, T-12) until T30 following insulin delivery (T0). Starting from T-12 leaded to positive AUC values.

### Food intake measurements

Food intake was measured following an overnight fasting in single-housed *Glp1r*^OBKORD^ RD mice and *Glp1r*^OBhM3Dq^ WD mice. SALB (10mg/kg) or CNO (3 mg/kg) were administered IP in *Glp1r*^OBKORD^ RD mice and *Glp1r*^OBhM3Dq^ WD mice respectively. Following drug administration, mice were given a pre-weighed amount of food (HFHS and/or Chow) with free access to water. Food was then weighed 1 hour after refeeding.

Mice were accustomed to daily manipulation 1 week before the test. Mice receiving a control treatment were administered an equivalent volume of saline. Comparison of treatments was done on a within-subject level (each mouse was its own control) and at least 1 week elapsed between each testing condition (drug or control treatment).

### Retrograde tracing

For retrograde tracing experiments, Alexa 488-conjugated cholera toxin B subunit (CTBgreen) was injected unilaterally at 0.05μL/min into the LH (0.3 μL; 1.5 μg/μL; A/P: -1.22 mm, M/L: -1.125 mm, D/V: -4.83 mm). Mice were culled 10 days after the injection. Brains were collected, post-fixed in 4°C PFA 4% overnight and cryoprotected in sucrose 30% for 2-3 days. Brains were then snap-frozen in isopentane and the OB was cut into 18 μm-thick sections afterwards, using a cryostat (Leica Biosystems). Brain slices were collected on microscope slides (SuperFrost Ultra Plus™ GOLD), mounted with mounting media (Fluoromount-G^TM^ with DAPI) and stored at -20°C until imaging. Image acquisition was performed with a confocal microscope (Zeiss LSM980 Airyscan2).

For experiments visualizing axonal projections from brainstem PPG neurons PPG-YFP mice and PPG-Cre mice were crossed, and their YFP- and Cre-positive offspring (1 male, 2 female) injected bilaterally with AAV2-Ef1a-DIO-tdTomato (0.25 μl/injection; UNC Vectorcore). Briefly, at 3 months of age these mice were anaesthetised with ketamine (50mg/kg; i.p.) and medetomidine (1mg/kg; i.p.) with meloxicam (5mg/kg, s.c.) given for peri-surgery analgesia. They were placed into a stereotaxic frame and their head was ventroflexed to enable access to the lower brainstem through blunt dissection of the neck muscles as described previously^7^. Injections were placed into the NTS (+/-0.50 mm M/L, +0.10 mm A/P, -0.35 mm D/V from obex) and into the IRT (+/-1.1mm M/L, +0.5mm A/P, -1.4mm D/V from obex) at 0.05μl/min. Three weeks later, mice were terminally anaesthetised with 100mg/kg pentobarbitol and transcardially perfused with PBS containing 4%PFA, brains were collected, post-fixed in 4% PFA at 4°C overnight and cryoprotected in 30% sucrose for 2-3 days. Brains were sectioned coronally at 30 μm thickness on a cryostat (Bright Instruments) and immunostained for YFP and tdTomato as described previously^7^. Brainstem, forebrain and OB sections were mounted on microscope slides, coverslipped and imaged on a Leica epifluorescence microscope.

### Noradrenaline levels

#### Drug administration and tissue sampling

Ex4 was injected in the OB of cannulated RD and WD mice, deprived of food for 5 hours. Mice were killed by decapitation and their brains were quickly removed and dissected on ice to separate the OB and the hypothalamus. Pancreas, gut, liver and epididymal adipose tissue were also dissected. Tissues were immediately deep frozen in liquid nitrogen, and stored at - 80°C until used. Each frozen tissue were weighed and dissolved in 200µL of lysing tissue solution (40 mg EDTA; HclO4 1mL of MilliQ water qs 100 mL) and then bead grinded for 2 minutes. The homogenates were centrifuged at 3000 rev/min for 30 min at 4°C, the supernatants were collected and centrifuged again at 16000 rev/min for 2 minutes. 20µL of the last supernatant was used for the HPLC-ED analysis.

#### High-performance Liquid Chromatography with Electrochemical Detection (HPL-EC)

NA measures were performed in a HPLC instrument (Shimadzu, LC20AD pump) equipped with a SiL20AC autosampler coupled with an electrochemical detector Waters 2465. NA separation was performed using an Ultrasphere column (Beckman, 150×4.6mm C18 5µM) equipped with two Phenomenex C18 filters in a security guard system. The mobile phase contained 150mM octane sulfonic acid, 8.2 mM ammonium acetate, 15% v/v methanol (pH=3.8, adjusted with glacial acetic acid) and was filtered (0.8mL/min, degassed before use) through a 0.2µM membrane filter. Elutes were detected at 700 mV (versus the reference electrode). The column and the detection cell were housed within the Faraday cage of the electrochemical detector (25.5°C). Samples were placed in the autosampler and kept at 4°C until the injection (20µl). Reference elution profiles were given by standard solutions of NA (50 ng/ml; Sigma-Aldrich) prepared in HCl 0.1N and stored at -20°C until use. Only pancreatic NA levels will be discussed in this thesis.

### Fluorescein-Trp25-Exendin-4 imaging

Fluorescein-Trp25-Exendin-4 was injected into the OB of cannulated RD and WD mice. After drug administration, an OGTT was performed as described before, until 90 minutes post-injection. At that moment, mice were transcardially perfused with PBS 1X followed by 4°C PFA 4% for 5 minutes. Brains were collected, post-fixed in 4°C PFA 4% overnight and cryoprotected in sucrose 30% for 2-3 days. Brains were then snap-frozen in isopentane and the OB was cut into 18 μm-thick sections afterwards, using a cryostat (Leica Biosystems). Brain slices were collected on microscope slides (SuperFrost Ultra Plus™ GOLD), mounted with mounting media (Fluoromount-G^TM^ with DAPI) and stored at -20°C until imaging. Image acquisition was performed with a confocal microscope (Zeiss LSM980 Airyscan2).

### LUXendin-645 imaging

To assess access of GLP-1R antagonists to the olfactory bulb from the circulation, three awake adult male PPG-YFP mice were injected subcutaneously with 30nM/kg LUXendin645 at a dose volume of 1ml/kg. After 2h survival the animals were terminally anaesthetised with 100mg/kg pentobarbitol and transcardially perfused with 30ml heparinised 0.1M PBS followed by 4% PFA in 0.1M PBS.

To assess access of GLP-1R antagonists from the ventricular system three adult (one male, two female) PPG-YFP mice were anaesthetised with ketamine (50mg/kg; i.p.) and medetomidine (1mg/kg; i.p.) and meloxicam (5mg/kg, s.c.) was given for peri-surgery analgesia. Mice were placed into a stereotaxic frame and a craniotomy was performed to allow unilateral microinjection of 5μl of 30μM into the left lateral ventricle (coordinates: +1 mm M/L, -0.5 mm A/P, -2.5 mm D/V from bregma). One hour after injection, still under anaesthesia, the animals were transcardially perfused with 30ml heparinised 0.1M PBS followed by 4% PFA in 0.1M PBS.

Brains were extracted and postfixed in 4% PFA overnight at 4°C. After cryoprotection with 30% sucrose in 0.1M PBS brains were sectioned coronally at 30μm thickness. Sections containing olfactory bulb, hypothalamus or lower brainstem were mounted on superfrost slides and coverslipped using Vectashield mounting medium. Subsequently, images were taken under a fluorescence microscope using excitation at 645 nm and emission at 664 nm for LUXendin-645 imaging, and 359 nm and 457 nm for DAPI.

### cFos immunohistochemistry

#### Tissue collection

Ex4 was injected into the OB of 5 cannulated RD and 5 WD mice cannulated mice. Mice receiving a control treatment (n=5 RD and 5 WD) were administered an equivalent volume of saline. After drug administration, an OGTT was performed as described before, until 90 minutes post-injection. Mice were then transcardially perfused with ice-cold PBS 1X followed by RT PFA 4% for 5 minutes. Brains were collected, post-fixed in PFA 4% at room temperature (RT) for 24h, under mild agitation. The following day, brains were transferred into PBS 1X on ice, under heavy agitation for 24h. Brains were cryoprotected in a 30% sucrose solution in PBS 1X for 2-3 days, at 4°C. Brains were snap-frozen in isopentane and the OB or LH was cut into 18 or 30 μm-thick sections respectively, using a cryostat (Leica Biosystems).

#### cFos staining in the OB

Slides were brought to RT and kept 5 min at 40°C in drying oven. Slides were outlined with a barrier pen and soaked 15 min in phosphate buffer (PB). Slides were washed 3 × 5 min in PB 0.2% Tx, incubated at RT overnight (22hrs) with 9F6 cFos rabbit antibody (1:1000; Cell Signalling Technology; #2250) in [PB 0.2% Tx + 1% donkey serum]. After 5 × 10 min PB washes, sections were incubated 2 h at RT with A568 donkey anti-rabbit secondary antibody (1:500; Invitrogen; A10042) in [PB 0.2% Tx + 1% donkey serum], 1% donkey serum. Slides were washed 4 × 10 min in PB, 1 × 5 mins in dH_2_0 and air dried (∼10 mins only). Slides were then mounted with Vectashield + DAPI (Vector labs H-1200).

#### cFos staining in the LH

On day 1, LH slices were washed 3×5 min in PBS 0.01M at RT and then incubated 30 min in [PBS 0.01M + 0.3% Triton + 50 mM Glycine + 2% NDS]. Subsequently, LH slices were incubated in guinea-pig anti-cFos (Synaptic Systems, 1:1000) diluted in [PBS 0.01M + 0.3% Triton + 10 mM Glycine + 0.1% H_2_O_2_]. O/N at RT – 475 µL/well. On day 2, slices were washed 3×5 min in PBS 0.01M at RT – 4 mL/well and incubated in donkey anti-guinea pig-AF647 (Jackson ImmunoResearch, 1:500) diluted in [PBS 0.01M + 0.3% Triton]. 2 h at RT – 475 µL/well. Next, 3 more washes of 5 min in PBS 0.01M at RT – 4 mL/well were performed and then LH slices were incubated in DAPI, diluted 1:20000 in PBS 0.01 M. 5 min at RT – 475 µL/well. One more wash 3x 5min in Tris-HCl 50mM (pH=7.5) at RT – 4 mL/well (12-well plate) was done before mounting the slices on gelatin coated slides (adding Prolong Gold antifade mounting medium, Invitrogen).

#### Imaging data capture and analysis

OB and LH sections processed for cFos in situ hybridization were imaged with a Slide Scanner (Nanozoomer 2.0HT, Hamamatsu, Japan). Image adjustments (brightness and contrast) as well as the generation of image montages were all performed using the open-source software ImageJ. For OB cFos image analysis, a custom-made pipeline (CellPofiler software) was used to automatically quantify cFos+ nuclei within the MCL. Total MCL area was measured using Phyton. Data was expressed as cFos+ nuclei/MCL area (mean from all animals). Poor quality slices or IHC stainings were excluded from the analysis. For LH cFos image analysis, the overlay of the atlas onto brain slices was performed using the ABBA plug-in for FIJI. cFos quantification was performed within the LH region and data was expressed as cFos+ nuclei/LH area (mean from all animals). 1 NaCl-injected and 1 Ex4-injected mouse were removed from the analysis due to uncomplete OB injection of Ex4.

### RNAScope

#### Tissue collection

*Glp1r*^OBhM3Dq^ mice (previously injected with viruses expressing hM3Dq-mCherry under the *Glp-1r* promoter control) were transcardially perfused with PBS 1X followed by 4°C PFA 4% for 5 minutes. Brains were collected on ice, post-fixed in 4°C PFA 4% overnight and cryoprotected in sucrose 30% for 2-3 days.

#### In situ hybridization of Glp1r mRNA

Sections from OB were processed using RNAscope *in situ* hybridization for *Glp1r* mRNA following the manufacturer’s protocol (RNAscope Multiplex Fluorescent Kit V2; Advanced Cell Diagnostics). 10μm-sections were cut on a cryostat (Leica Biosystems) and collected on Superfrost Plus slides, then air-dried at room temperature for 1h followed by incubation at 40°C for 5mins. Slides were then incubated in rising concentrations of ethanol (50%, 75%, 95%, 100%; each 5mins) and treated with H_2_O_2_ for 10mins at room temperature. Following incubation in protease III for 30mins at 40°C, sections were incubated in a probe recognizing murine mRNA for *Glp1r* (ACD, Cat No. 418851-C2) at 40°C for two hours. Following amplification according to the manufacturer’s protocol, sections were incubated in HRP-C2 at 40°C for 15mins followed by Opal650 (Akoya) in TSAP diluent for 30mins at 40°C. Following RNAscope, slides underwent immunolabelling for mCherry. Slides were incubated for 18 hours at room temperature in anti-dsRed antibody (TaKaRa, 1:2000) in 0.1M PB containing 0.3% Triton X-100 and 1% donkey serum. Slides were incubated in AlexaFluor 488-conjugated anti-rabbit secondary antibody in 0.1M PB containing 0.3% Triton X-100 and 1% donkey serum for two hours at room temperature and immediately covered using Vectashield mounting media. OB sections were imaged with a Leica TCS SP8 confocal microscope at x20. Image adjustments (brightness and contrast) as well as the generation of image montages were all performed using the open-source software Fiji.

### GLP-1 content on OB slices

Ten RD male mice and 10 WD male mice (17 weeks-old) were transcardially perfused with ice-cold NMDG saline buffer. The brain was harvested and the olfactory bulb dissected and immediately sectioned in 12 slices of 150µm using a vibratom. Slices were then transferred to NMDG at 34°C for ∼12 min and divided into different wells (∼1 bulb per well). Slices were then incubated for 10 minutes at 34°C (monitoring of bubbling, keeping slices in motion using a pipette) in wells containing 600µL of aCSF at 34°C (including 1mM or 5mM glucose) under fine bubble carbogen perfusion (1Hz). Protease inhibitors (DPP4i at 24µL/600µL) and Aprotinin (9µL/600µL) were added before transfer of slices into the wells. Slices were then transferred to wells filled with 600 µL of ACSF (+100mM KCl) with DPP4 inhibitors (24µL/600µL) and Aprotinin (9µL/600µL) and incubated for 20 min at 34°C (monitoring of bubbling, keeping slices in motion using a pipette). The 1^st^ incubation wells (ACSF + inhibitors) were aspirated (bottom of the well (∼500µL)) with a pipette. Supernatant samples were named “T0” and stored at -80 until dosage. Next, the 2^nd^ incubation wells (ACSF + inhibitors + KCl) were aspirated. Supernatant samples were named “T20” and stored at -80 until dosage. OB slices were recollected apart. GLP-1 content in the supernatant of T0 and T20 samples was assessed using a Glucagon-Like Peptide-1 (GLP-1) Total ELISA KIT (96-Well Plate Cat. # EZGLP1T-36K, EZGLP1T-36BK, Millipore).

### Real time quantitative PCR

All animals were sacrificed by cervical dislocation. The brain was harvested and the olfactory bulb and hypothalamus were dissected and immediately deep frozen in liquid nitrogen and stored at -80°C. Total RNA was isolated using Rneasy Lipid Tissue mini kit (Qiagen). Real-time quantitative polymerase chain reaction (RT-qPCR) was carried out in a LightCycler 480 detection system (Roche) using the LightCycler FastStart DNA Master plus SYBR Green I kit (Roche). The primers were derived from mouse sequences (Table S1). We targeted *Glp1r*, *Pcsk1*, *Dpp4* and *Ppg* normalized against the mean of two reference house-keeping genes: *Rpl19* and *Tbp*.

### Quantification and statistical analysis

Data in representative are given as means ± standard error of the mean (SEM). Statistical analysis was performed using Student’s t test (to compare the means of specifically two groups) or ANOVA (to analyze the difference between the means of more than two groups) after normality was assessed by a Shapiro-Wilk test. One-way ANOVA or two-way ANOVA tests were selected when data was collected about one or two independent variables, respectively. ANOVA tests were followed by two-by-two comparisons using Bonferroni unless otherwise stated (GraphPad Software, La Jolla, CA, USA). Differences were considered significant at p < 0.05.

