## Supplementary Figures for "A neuronal circuit driven by GLP-1 in the olfactory bulb regulates insulin secretion"

Figure S1

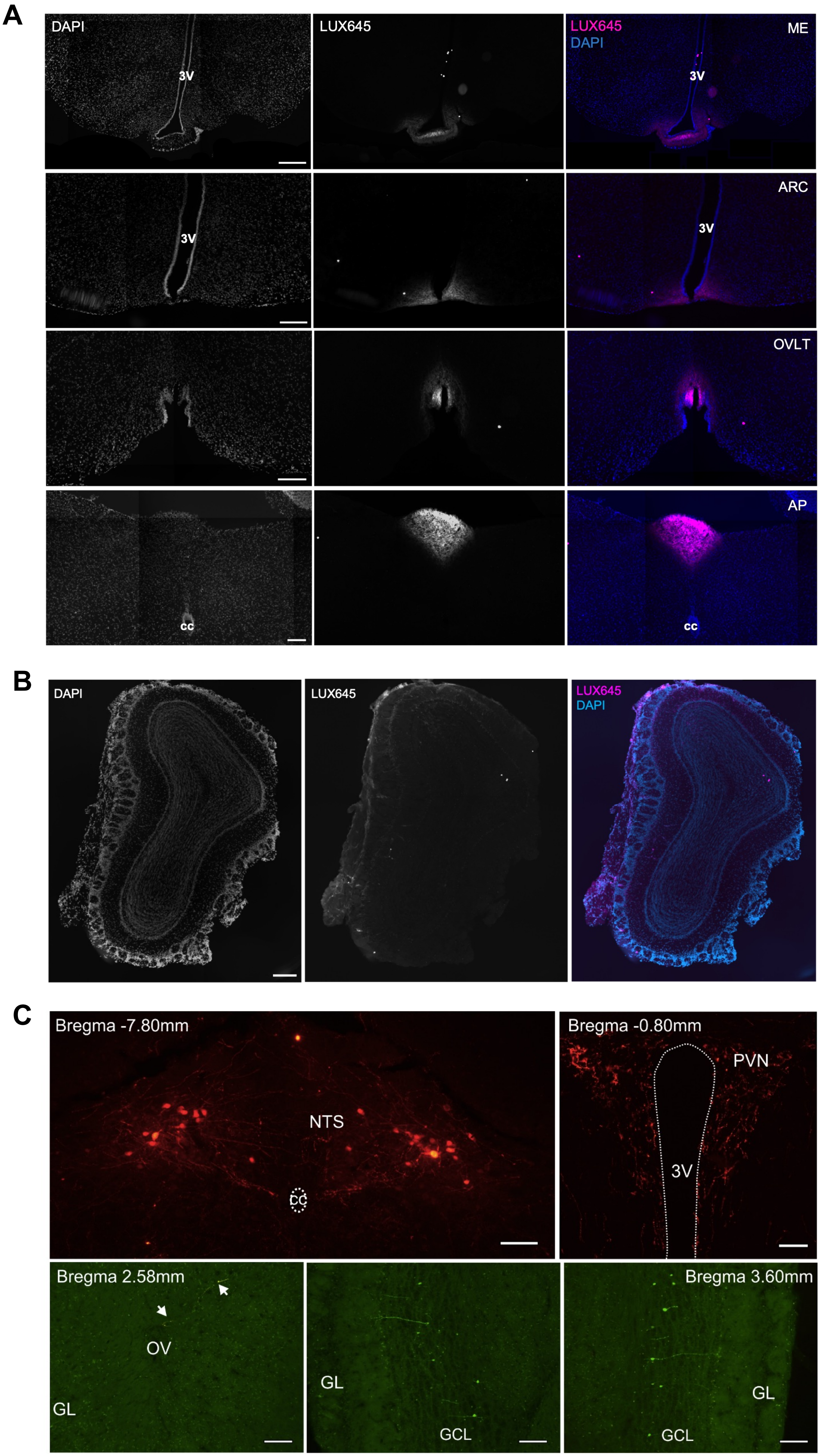
A
DAPI
LUX645
LUX645
DAPI
ME
3V
3V
ARC
3V
3V
OVLT
AP
cc
cc
B
DAPI
LUX645
LUX645
DAPI
C
Bregma -7.80mm
NTS
CC
Bregma -0.80mm
PVN
3V
Bregma 2.58mm
OV
GL
GL
GCL
Bregma 3.60mm
GL
GCL

**Figure S1. Deciphering the GLP-1 neuronal circuit in the OB**

(A) Coronal brain sections of a mouse IP-injected with LUXendin 645 (LUX645). From top to bottom: images showing LUX645 staining in the median eminence (ME), arcuate nucleus (ARC), organum vasculosum laminae terminalis (OVLT) and area postrema (AP). Scale bar: 200 µm.

(B) Photomicrograph of representative OB coronal sections of a mouse icv-injected with LUXendin 645 (LUX645). Scale bar: 200 µm.

(C) Targeting of the $PPG^{NTS}$ neurons with an AAV expressing tdTomato (top left). PVN axons of $PPG^{NTS}$ neurons targeted with an AAV expressing tdTomato (top right). Virally transduced fibers close to the olfactory ventricle at Bregma 2.58mm (bottom left). Overlays of YFP and tdTomato fluorescence in OB slices showing PPG neurons in the GCL (bottom middle and right). Scale bar: NTS, nucleus tractus solitarii; PVN, paraventricular nucleus of the hypothalamus; GL, glomerular layer; OV, olfactory ventricle; GCL, granular cell layer; 100 µm. Related to Figure 1.

#### Figure S2

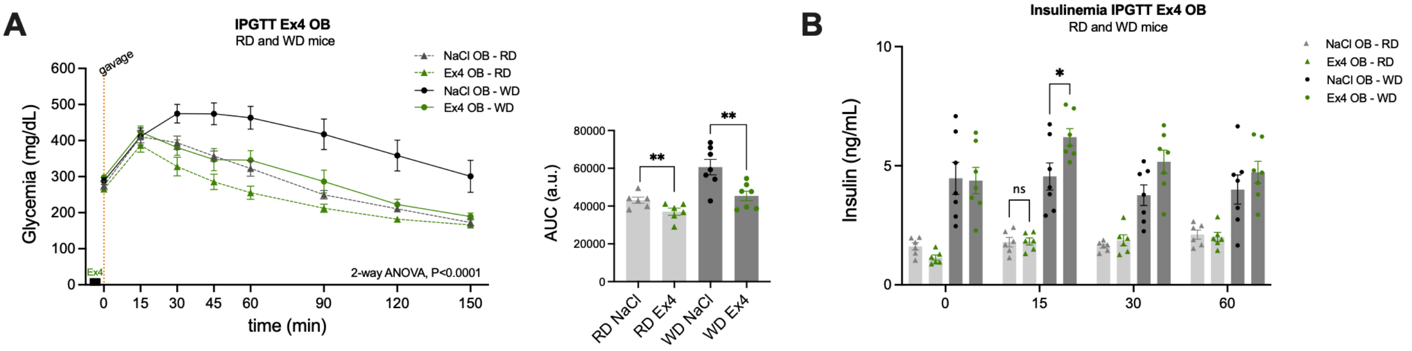


**Figure S2. IPGTT and insulinemia after the injection of Ex4 in the OB.**

(A) IPGTT tests combined with preceding OB injections of Ex4 (n=6 RD and 7 WD) in OB-cannulated RD and WD mice after 20 weeks under WD. Black squares on X axis indicate drug delivery before the glucose challenge (T0) (left). AUC of glycaemia (right). OGTT curves are analyzed using Repeated Measures (RM) two-way ANOVA followed by Bonferroni post-hoc test; AUCs are analyzed using Student's t-test followed by Bonferroni post-hoc test.

(B) Plasma insulin levels of OB-cannulated RD and WD mice measured during Ex4 OB-injected IPGTT. Data are analyzed using two-way ANOVA followed by Bonferroni post-hoc test (n=6 RD and 7 WD).

Data are given as mean ± SEM. *p <0.05; **p < 0.01; ***p < 0.001; ns, not significant. Related to Figure 2.

### Figure S3

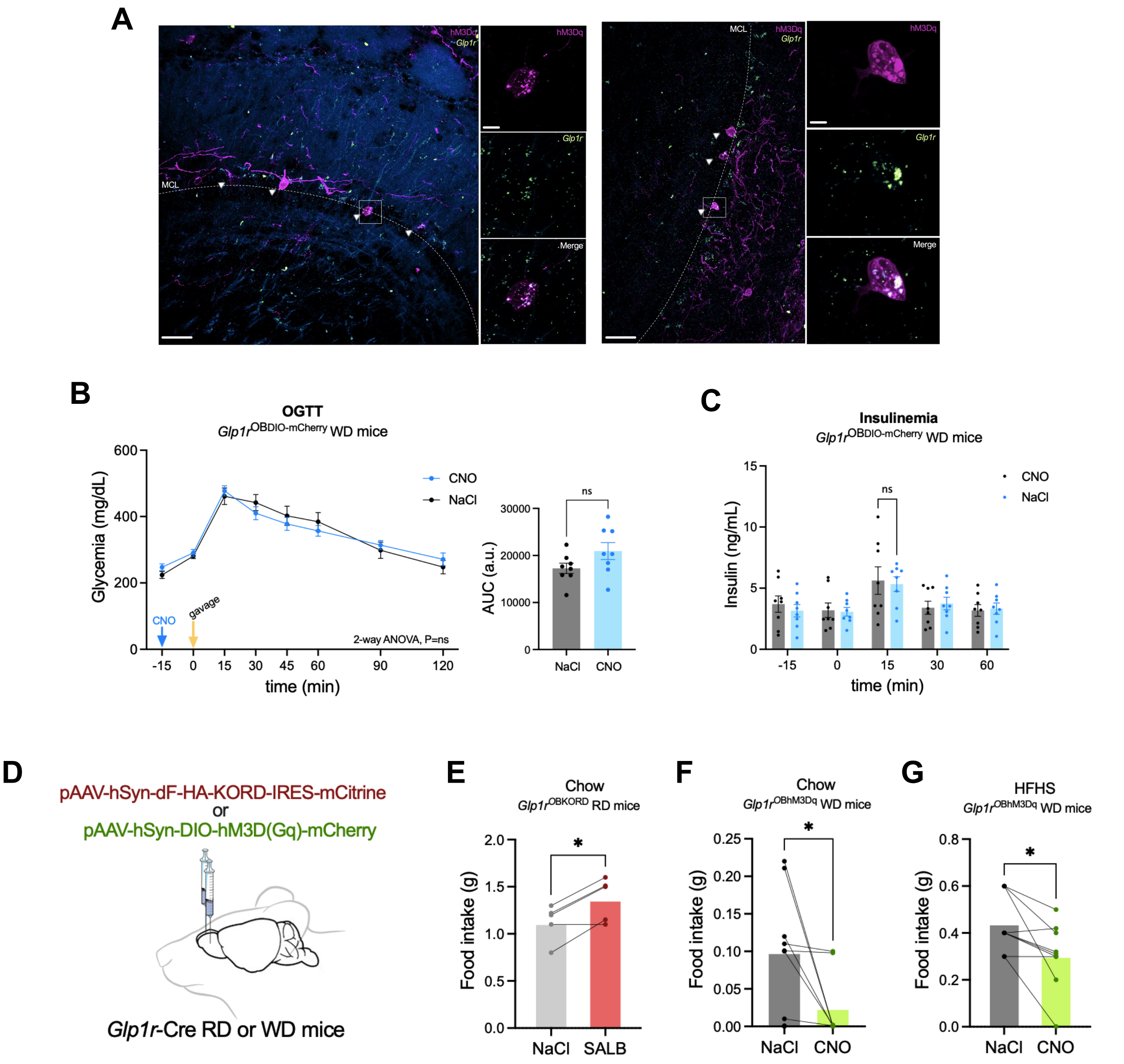


**Figure S3. Further characterization of the DREADD mouse model.**

(A) Representative images of RNAscope for *Glp-1r* in hM3Dq-mCherry-injected *Glp1r* Cre mice. Scale bar = 50 µm and 20 µm (higher magnification images). Fluorescence of mCherry is expressed in magenta and *Glp-1r* mRNA is expressed in green. Related to Figure 3.

(B) OGTT tests preceded by an IP injection of CNO in *Glp1r*$^{OBDIO\text{-}mcherry}$ WD mice after 20 weeks under WD (26-week-old, n=8). The blue arrow indicates the CNO injection before the glucose gavage (T0) (left). AUC of glycaemia (right). OGTT data are analyzed using RM two-way ANOVA followed by Bonferroni post-hoc test and AUCs by using paired Student's t-test.

(C) Plasma insulin levels of *Glp1r*$^{OBDIO\text{-}mcherry}$ WD mice after CNO IP injection (RM two-way ANOVA followed by Bonferroni post-hoc test; n=8). Data are given as mean ± SEM. *p <0.05; **p < 0.01; ***p < 0.001; ns, not significant. Related to Figure 3.

(D) Schematic illustration of viral delivery in the OB of *Glp1r*-Cre RD and WD mice.

(E-G) Food intake in *Glp1r*$^{OBKORD}$ RD mice (n=5) fed a Chow diet (E) and in *Glp1r*$^{OBhM3Dq}$ WD mice (n=9) fed both a Chow (F) and a HFHS (G) diet 1h after IP injection of SALB and CNO, respectively. Data are analyzed using Paired Student's t-test. Data are given as mean ± SEM. *p <0.05; **p < 0.01; ***p < 0.001; ns, not significant. Related to Figure 3.

### Figure S4

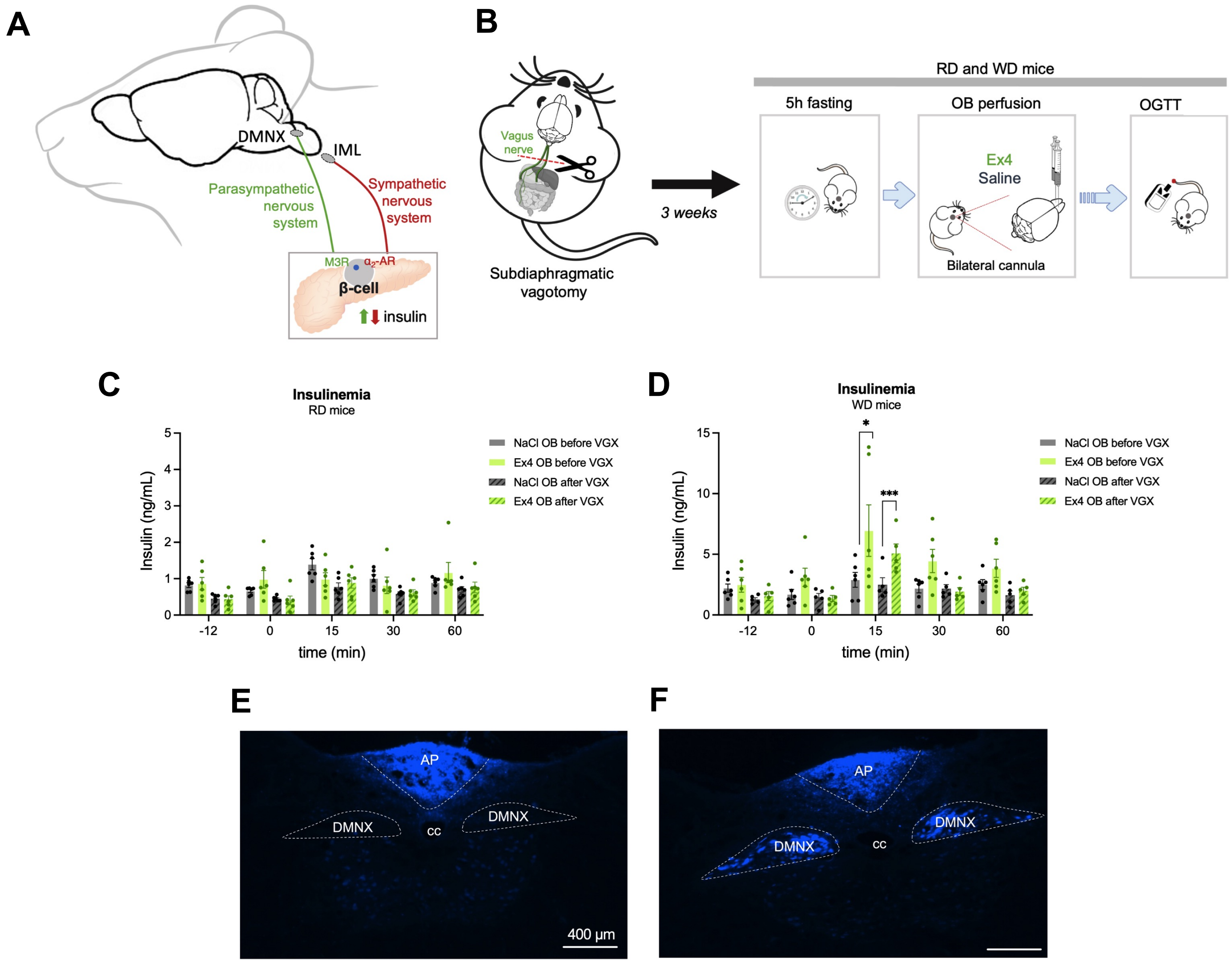


**Figure S4. The enhancing effect of Ex4 injection in the OB on glucose tolerance and insulin secretion persists after vagotomy.**

(A) Parasympathetic and sympathetic control of insulin secretion.

(B) Schematic illustration of bilateral OB injections of Ex4 followed by an OGTT in OB-cannulated vagotomized WD and OB-cannulated vagotomized RD mice.

(C) Plasma insulin levels of OB-cannulated RD during OGTT tests preceded with OB-injected Ex4 before and after vagotomy (RM two-way ANOVA followed by Bonferroni post-hoc test, n=6).

(D) Plasma insulin levels of OB-cannulated WD during OGTT tests preceded with OB-injected Ex4 before and after vagotomy (two-way ANOVA followed by Bonferroni post-hoc test, n=5-6/group).

(E-F) Representative photomicrographs of fluorogold staining in the DMNX. Absence of fluorescence in vagomotized mice (left image) or fluorescence labeling in sham mice (right mice). Images taken with a fluorescence microscope (Scale bar = 400 µm). AP, area postrema; DMNX, dorsal motor nucleus of the vagus; cc, corpus callosum.

Data are given as mean ± SEM. *p <0.05; **p < 0.01; ***p < 0.001; ns, not significant. Related to Figure 4.
