## Supplementary material for "A neuronal circuit driven by GLP-1 in the olfactory bulb regulates insulin secretion": Suplementary Table

**Table S1. List of primers used for RT-qPCR of *Glp1r*, *Psck1*, *Dpp4* and *Ppg* in the OB of RD and WD mice.**

| Gene | Forward | Reverse |
|---|---|---|
| *Glp1r* | CGGCGTCAACTTTCTTATCTTCA | GGGCGTGTTCGTCCATCA |
| *Pcsk1* | TGGAGTTGCATATAATTCCAAAGTT | AGCCTCAATGGCATCAGTTAC |
| *Dpp4* | CGGTATCATTTAGTAAAGAGGCAAA | GTAGAGTGTAGAGGGGCAGACC |
| *Ppg* | TACACCTGTTCGCAGCTCAG | TTGCACCAGCATTATAAGCAA |
| *Rpl19* | TCAAAAACAAGCGGATTCTCA | GCGTGCTTCCTTGGTCTTAG |
| *Tbp* | GGGTTTTCCAGCTAAGTTCTTG | AGCACAAGGCCTTCTAACCTT |
